# Slowing down an analyte in a nanopore

**DOI:** 10.1101/2021.01.11.426231

**Authors:** G. Sampath

**Affiliations:** Unaffiliated

**Keywords:** single molecule, nucleotides, amino acids, DNA, peptides, Fokker-Planck equation

## Abstract

An unresolved problem in nanopore sensing is the high translocation speed (∼10-100 monomers/μs) of an analyte (nucleotide, DNA, amino acid (AA), peptide) through the pore. Here a method based on reversing the pore voltage and changing the solution pH is described. A simplified Fokker-Planck model shows mean translocation times of 1-10 ms in a nanopore of length 10 nm. Simulations show that a positive-negative voltage profile can trap an analyte for ∼1 ms. This method can be used for free nucleotides, single AAs, oligonucleotides, and oligopeptides. Its applicability to existing nanopore sensing and sequencing techniques and implementation issues are discussed.

## 1. Introduction

In recent years nanopores [1,2] have emerged as a single-molecule alternative to established methods of polymer sequencing and analysis. Most reports of nanopore studies of an analyte (DNA/RNA strand, oligonucleotide, protein/peptide, free nucleotide, free amino acid (AA)) are usually focused on one of four analytical functions: detection, identification, sequencing, and chemical analysis (of chemical interactions, structure, and post-translational and other modifications). Nanopores are on the way to becoming a viable alternative for DNA sequencing at the single molecule level [1,2]; nanopore protein sequencing is yet to show a similar degree of success [3].

Nanopore analysis [1] is based on the use of an electrolytic cell (e-cell) with *cis* and *trans* chambers separated by a thin membrane with a nano-diameter pore; a polymer translocates through the pore via diffusion and electrophoresis (and, in some cases, electroosmotic (EO) flow [4-7]). Sensing may be based on measuring the resulting current blockade [1,2], optical detection of fluorophore labels attached to the analyte [8], or transverse current measurements lateral to the pore [9].

Broadly there are two approaches to DNA sequencing: ‘strand sequencing’ [1,2] and ‘exonuclease sequencing’ [10]. The most successful strand sequencing method thus far is ‘sequencing by synthesis’ (SBS) [11,12]. With exonuclease sequencing, base calling is easier, at least in principle; errors can be reduced by using a modified e-cell with two pores in tandem [13,14]. These methods are based on measuring blockade currents; in contrast optical methods use fluorescent labels [8,15] and require much less precision. Alternatively sequencing can be based on transverse currents [9] that are 1 to 2 orders of magnitude higher than longitudinal pore currents.

Nanopore protein sequencing is considerably more difficult than DNA sequencing in large part because there are 20 AAs to identify, as opposed to 4 bases in DNA. A variety of methods are known [16-19], some theoretical, others practical but yet to be implemented fully; for recent reviews see [20-21]. The nanopore approach faces a major difficulty. The analyte translocates through the pore at a high rate, about 10-100 monomers per microsecond, which is beyond the capability of the detector. A wide-ranging solution has remained elusive; most efforts are limited to one type of analyte, usually single DNA strands. Two reviews separated by a decade show that progress is far from complete [22,23]. Examples of slowdown methods include [24-28]. Most of these are focused on DNA strands; a few (such as [29]) have looked at protein/peptide slowdown as well. EO can in some situations dominate electrophoresis and even reverse the direction of translocation, thus induced EO has been proposed [4] and used as an explicit slowdown mechanism [6].

### The present work

This letter introduces a slowdown method that can be used with a variety of analyte types: single nucleotides, free AAs, oligonucleotides, and peptides. A combination of theory, modeling, and simulation is used to show that residence/translocation times of ∼1 ms in a nanopore ∼10 nm long can be achieved. The method (a simpler version of which was mooted earlier in the Supplement to [13]) runs counter to the approaches in [24-29]; it is based on *reversing the voltage across the pore* and *increasing the mobility of some analytes (by changing the solution pH)*. A simplified Fokker-Planck (F-P) model of analyte translocation shows significant translocation time increases, up from tens of nanoseconds to the millisecond range, for the four nucleotides (dAMP, dTMP, dCMP, dGMP), the 20 AAs, short homonucleotides and heteronucleotides, and short homopeptides and heteropeptides. Simulations more realistic than the model show that a bi-level positive-negative voltage profile across the nanopore can yield translocation times of ∼1 ms, which can be detected with a bandwidth of 1-10 Khz.

In the model presented here EO (which does not affect all analytes equally) is excluded for two reasons: 1) to simplify the model; and 2) its effect can be nullified so that translocation can be controlled through the applied voltage. The role of EO in nanopore analysis is further discussed in Section 6 of the Supplement.

In the e-cell of Figure 1A, the electrophoretic velocity of an analyte is determined by its mobility, which is proportional to the analyte’s charge. dAMP, dTMP, dCMP, and dGMP carry roughly the same amount of negative charge and go from *cis* to *trans*. The 20 amino acids in

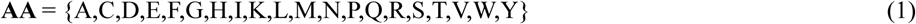

carry a charge equal to C_mult_q, where 0 ≤ |C_mult_| ≤ 2 and q is the (absolute) electron charge. C_mult_ depends on the pH level of the solution, it can be calculated with the Henderson-Hasselbalch (H-H) equation [30]. To reduce the number of AAs to consider when devising methods to slow them down, they are divided into equivalence classes. Two such are:

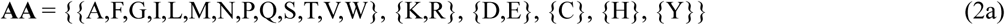

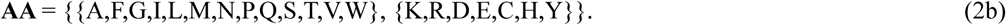

Two AAs are in the same class if their maximum absolute mobilities are roughly the same (or at least of the same order). To calculate the mobilities of oligonucleotides and oligopeptides, these polymers are modeled as rigid rods. The details are given in the Supplement (Section S-1), where detailed calculations of C_mult_ for free AAs and oligopeptides are presented (see Equations S-1 through S-11 therein).

**Figure 1.**
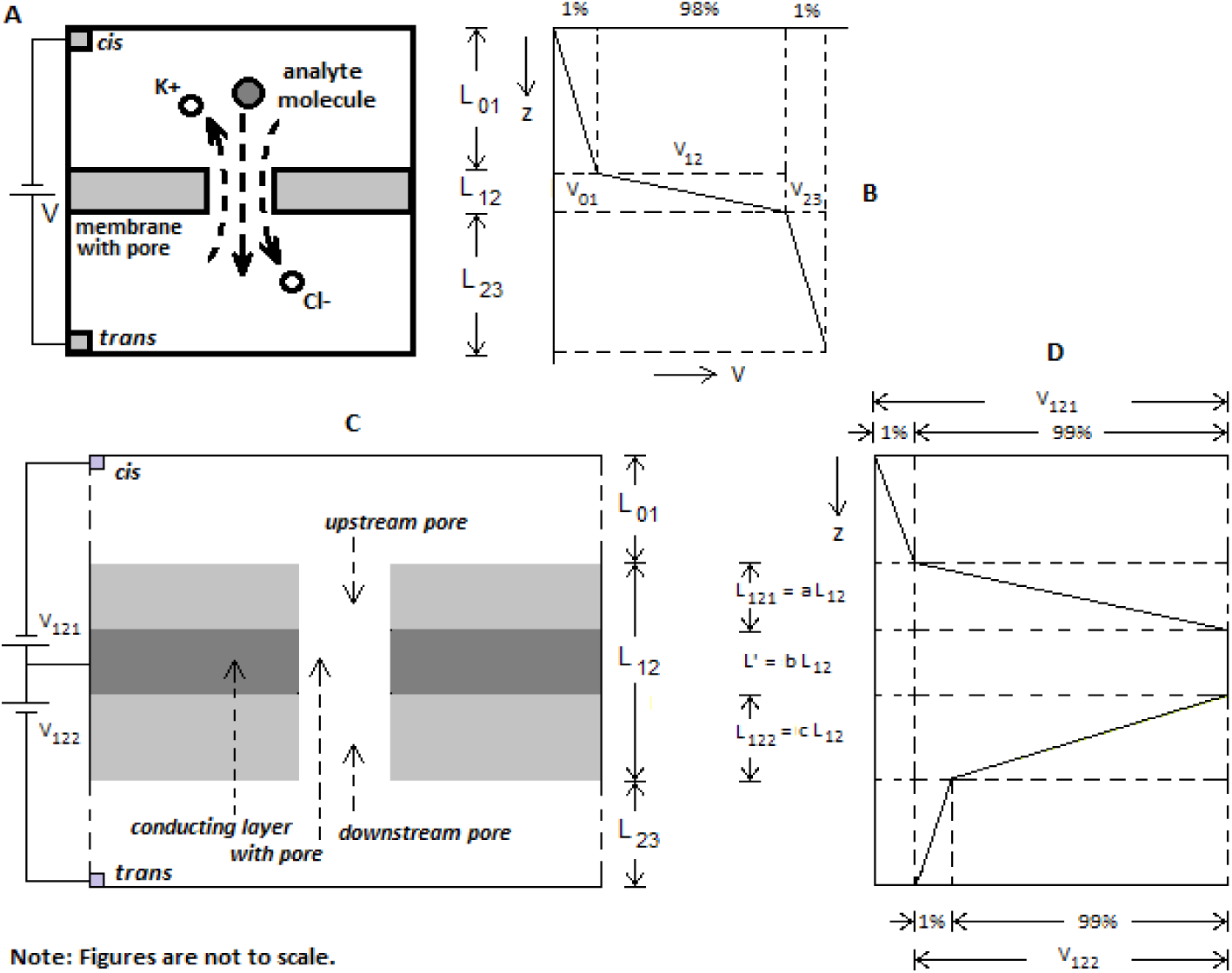
Schematic of electrolytic cell (e-cell); membrane containing nanopore separates *cis* and *trans* chambers containing salt (KCl) solution. (A) Ionic current flow in e-cell is due to K^+^ and Cl^-^ ions; analyte translocating through pore causes reduction in open pore current; (B) 98% of voltage V between *cis* and *trans* drops across pore; (C) Two pores in tandem with different voltages across each and intervening conducting membrane layer; (D) Positive voltage V_121_ across upstream pore of length L_121_ = aL_12_; 99% drops across pore, remaining 1% across *cis*; conducting membrane layer at constant potential over thickness bL_12_; negative voltage V_122_ across downstream pore of length L_122_ = cL_12_, 99% drops across pore, remaining 1% across *trans*. In simulations L_12_ = 11 nm, a = c = 5/11, b = 1/11. Voltages V_121_ and V_122_ are varied to give different profiles so analyte can be trapped for long durations; their values depend on analyte and pH of solution, see Figure 2.

An F-P equation is used to model analyte motion in the e-cell in Figure 1A and compute the dwell/translocation time of an analyte in/through the pore. Let T be the time of translocation of an analyte through the pore and E(T) its mean. Let D be the diffusion constant, μ the analyte mobility, and v_z_ = μV_12_/L the drift velocity through the pore. With α = v_z_L/D = μV_12_/D, V_12_ = V, and L_12_ = L, it can be shown that

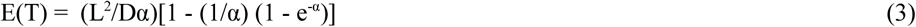

The following three cases can be considered:

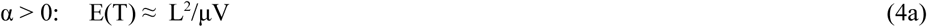

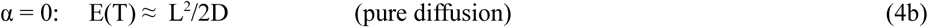

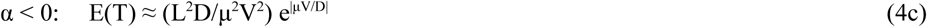

For a summary of the mathematical analysis leading to Equation 3 see the Supplement (Section S-3); the details can be found in the Supplement to [13].

Equation 4c shows that *reversing the voltage V*_*12*_ *across the pore and increasing the mobility of an analyte (by changing the pH of the solution) can significantly increase the translocation time*. Table 1 shows translocation times in a nanopore of length 10 nm (at different pH values) for: 1) dAMP, which represents all five nucleotides; 2) Arginine, which represents the AA subset {K, R, D, E} (see Equation 2b); 3) Cysteine, Histidine, and Tyrosine individually (corresponding to {C}, {H}, and {Y} in Equation 2a); 4) Alanine, which represents {A,F,G,I,L,M,N,P,Q,S,T,V,W} (see Equation 2b) at two different values of pH; 5) the homonucleotide AAAAAAAAAA, where A represents dAMP); 6) CCAGTTTATA, a random heteronucleotide; 7) the homopeptide RRRRRRRRRR; 8) HWVEDVDLTP, a random heteropeptide with both charged and uncharged residues; 9) IVHSMSWALP, a random heteropeptide with one charged residue (H); 10) IVFSMSWALP, same as 9 but with the uncharged residue F replacing the charged residue H, at two different pH values; and 11) the peptide in 10 with a positively charged header RRRRR. This is a comprehensive mix that covers all cases of interest.

**Table 1.**
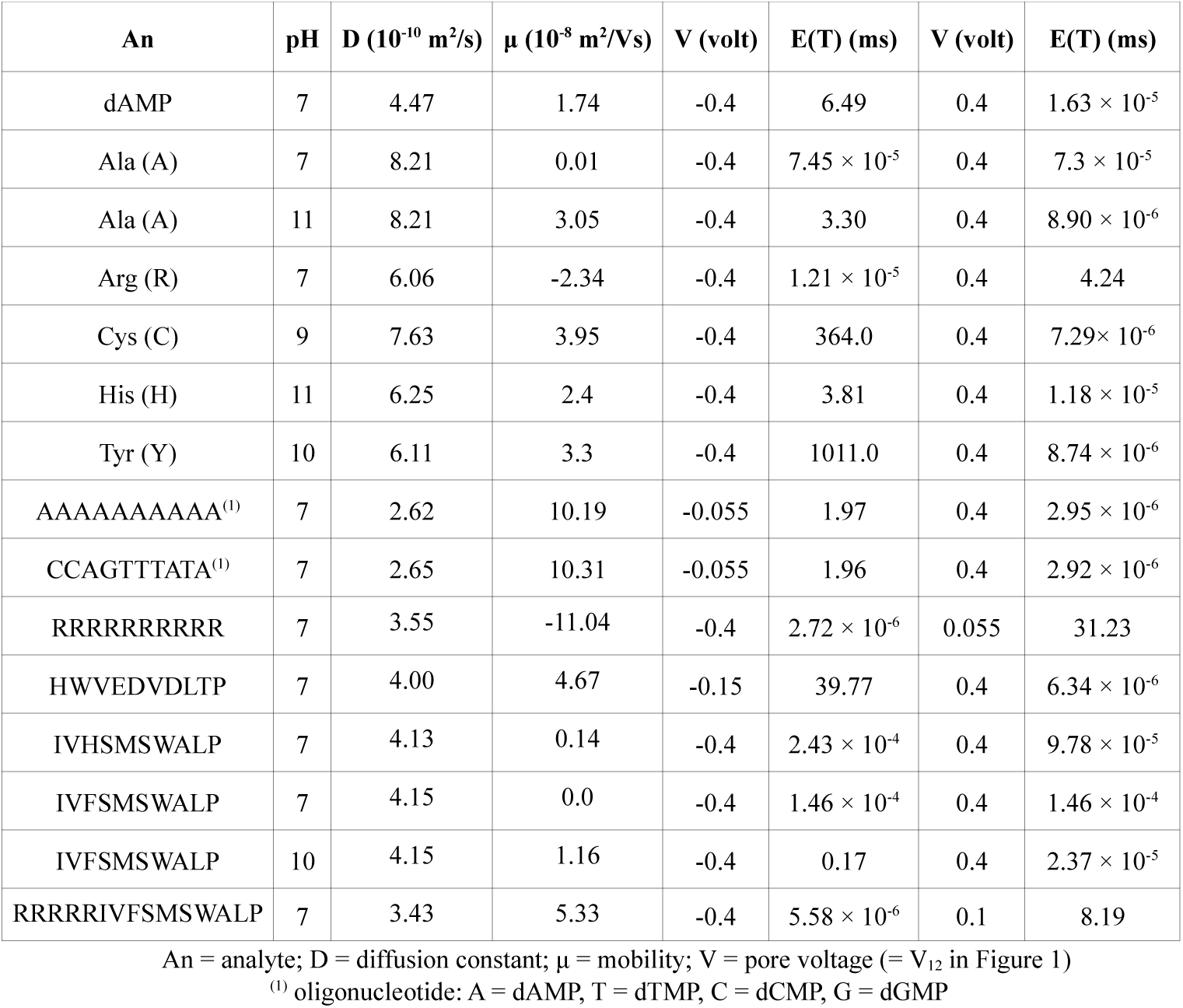
Mean translocation times E(T) for 15 analytes in tandem pore of length 11 nm (L_12_ in Figure 1A)

In what follows, the analyte is assumed to be negatively charged. The development can be extended to positively charged analytes with a systematic change of sign.

The high translocation times in Table 1 are mainly due to the front end of the pore being considered a reflector in the F-P model, which views the pore in isolation. With the tandem pore structure of Figure 1C, no such reflective boundary exists for the upstream pore, which is open at both ends. However, it can be approximated with the tandem pore structure in Figure 1D. Thus with a positive voltage applied to the upstream pore the analyte moves toward the downstream pore; effectively *the upstream pore with the positive voltage is acting like a reflecting boundary for the downstream pore with the negative voltage*. The values of the positive and negative voltages required to trap the analyte inside the structure for a long time can be obtained approximately by simulation (as has been done here) and more accurately by experiment.

This analysis neglects the role of electroosmosis (EO), which occurs due to water molecules flowing in the reverse direction (to the electric field) in the presence of charges on the inner surface of the pore. This effect, which is a countervailing force to the normal flow due to diffusion and electrophoresis, could in extreme cases reverse the direction of movement of the analyte. In the model presented here EO is excluded to simplify the model and because EO effects are virtually absent if the inner pore surface is coated with a charge-neutralizing lipid coat [31] (thus allowing analyte translocation control to be exercised primarily through electrophoresis).

A rough estimate of the expected translocation time E_12_(T) through the tandem pore structure in Figure 1C can be obtained by adding the times for the three segments of the pore (Equation 4). However because of ever-present diffusion this estimate can be considerably off as the analyte goes through four discontinuities while translocating through the tandem pore structure of Figure 1C. One way to get better estimates is to formulate a discontinuous F-P equation for the structure in Figure 1C and solve it numerically. Here an alternative approach based on simulation is taken. The simulation is based only on diffusion and electrophoresis; EO is excluded.

The slowdown process described above was simulated by assuming an analyte molecule to be a dimensionless particle. The procedure is similar to that in [14]; the details are available in the Supplement. Of the 15 analytes in Table 1, 11 were used in the simulations. Because of the bi-level voltage profile an analyte may oscillate between *cis* and the tandem pore, stay for extended times inside, or translocate fully into *trans*. In the absence of full translocation the longest dwell time of the analyte during the simulation run is used. Figure 2 shows the increases in translocation times achieved with different bi-level voltage profiles. Notice how the time for Alanine jumps by ∼4 orders of magnitude when the pH goes from 7 to 12 in Figure 2A. Notice also the much smaller negative voltage required to slow down AAAAAAAAAA, CCAGTTTATA, and RRRRRRRRRR, which is sufficient to give a residence time of >2 ms (Figure 2B). In all cases, with more negative voltages the analyte can be trapped for an indefinite length of time.

**Figure 2.**
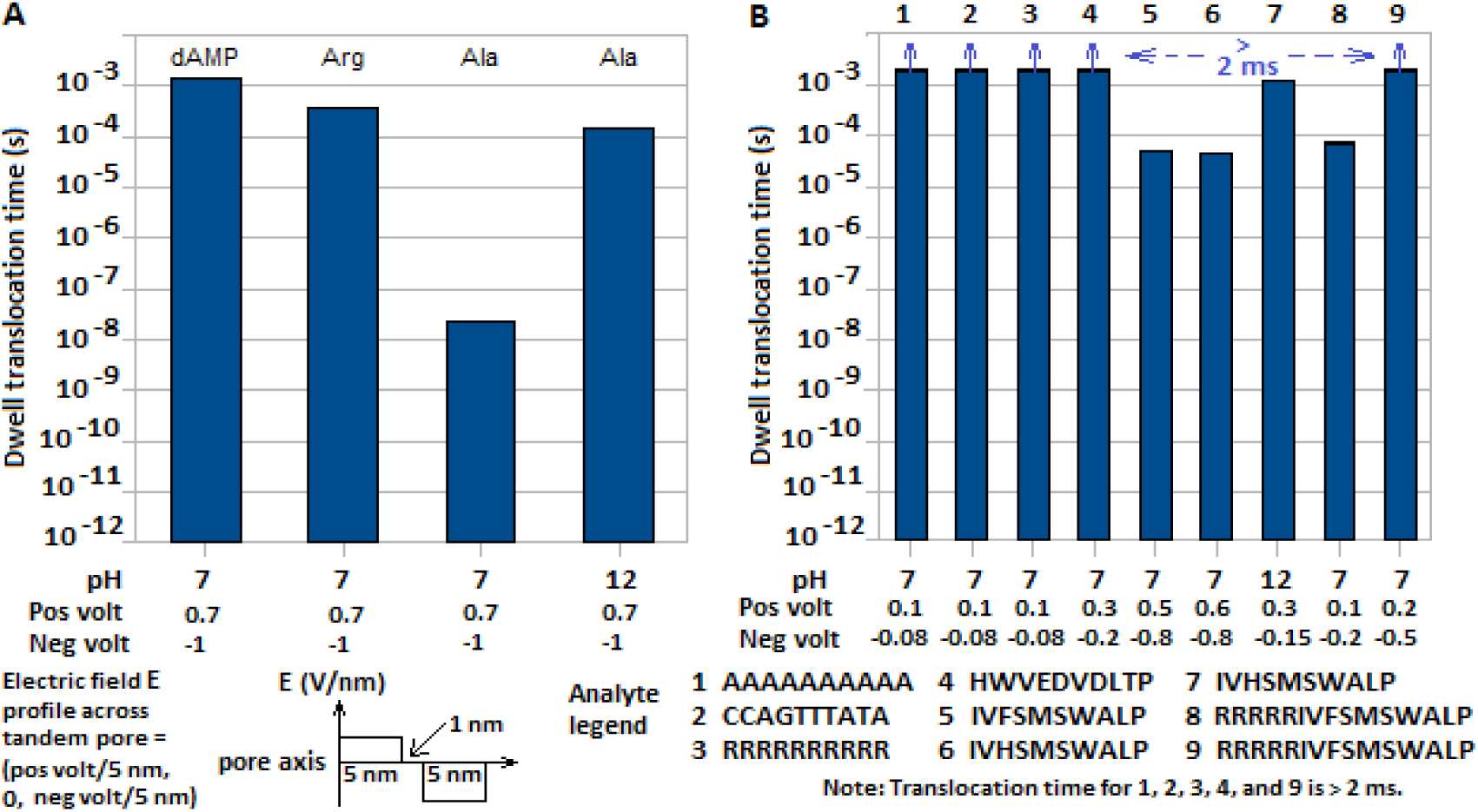
Mean translocation times of nine analytes for different voltage profiles over tandem pore structure in Figure 1C. Profiles shown are of electrical field dV/dz, which is constant across each segment of tandem structure. Lengths on z-axis correspond to tandem pore segment thicknesses; L_121_ = aL_12_, L’ = bL_12_, L_122_ = cL_12_ in Figure 1; a = c = 5/11, b = 1/11, L_12_ = L = 11 nm used in simulations. Electric field (V/nm) in profile is the triple (positive field across upstream pore, conducting layer field (= zero), negative field across downstream pore).

Time data correspond to larger of dwell time and translocation time through tandem structure. (A) Translocation times for dAMP, Arg, and Ala with different electrical field profiles and pH values; (B) Translocation times for two oligonucleotides and six oligopeptides with different electrical field profiles and pH values. (Pos volt and Neg volt correspond respectively to V_121_ and V_122_ in Figure 1.)

An unintended consequence of the bi-level voltage profile is a decrease in the pore ionic current. Its magnitude can be estimated from the translocation/dwell times of ions in the pore. Since the current delivered by an ion from *cis* to *trans* is inversely proportional to the translocation time through the pore that separates the two chambers the result is a steep fall in the current to ∼10% of that without the opposing voltage. This is elaborated in the Supplement; see Table S-7 therein. However, if the negative segment voltage is low (see, for example, the profiles in Figure 2B for AAAAAAAAAA, CCAGTTTATA, and RRRRRRRRRR) this reduction is negligible. Even with a peptide that carries a low charge, it can be invested with a high charge by attaching a short homopeptide made up of charged residues. For example RRRRR can be attached to the uncharged heteropeptide IVFSMWALP; notice the resulting increase in the translocation time to > 2 ms. However as noted earlier (and discussed further in Section 4 below) sensing need not be based on blockade currents, optical methods [8,15] or transverse currents [9], which are not dependent on the pore current, may be used instead.

## 4. Discussion

1. The method does not require any special devices or chemicals. The structure in Figure 1C can be fabricated as a stack of three layers: two synthetic silicon-based membranes and a conducting membrane in between, with graphene or aluminum for the latter. Fabrication of nanopore-based structures is reviewed in [32].
2. Two previous theoretical studies [25,26] have considered the use of multi-level voltage profiles for analyte slowdown. In [25] a multilayer structure with alternating insulator and metal layers is proposed for DNA sequencing; it requires a ∼250 MHz driving signal. This does not take into account the effect of high frequency noise in the pore [33], which makes measurement of blockade current differences difficult. In [26] a pore in a 2- or 3-layer graphene membrane is studied with molecular dynamics (MD) simulations. The potential of the membrane is switched from positive to negative after the strand has partially passed through the pore. The total MD simulation time is given as 860 ns. It is not clear if the results can be extrapolated to the millisecond range, where useful blockade current measurements can be made without the deleterious effects of noise.
3. The slowdown method given here may be well suited for use with exonuclease sequencing of DNA [10].
4. In [15] a fluorophore-labeled molecule passes from the unlit *cis* side of an e-cell to the lighted *trans* side where it is detected. If the pore is replaced with a channel open on one side along its length it may be possible to optically record the analyte’s passage through the channel.
5. With AAs, since different pH values may be required to achieve slowdown, they may need to be probed in different e-cells in series. A method to separate AAs from a sample by routing 20 copies of the AA to 20 channels, each containing a different transfer RNA (tRNA), is described in [34]. In exactly one of these channels the corresponding (cognate) tRNA gets ‘charged’ with (that is, bound to) the AA. Following this the bound AA is released through the addition of NaOH, and the freed AA enters an e-cell, where its presence is detected from the mere occurrence of a blockade, it is not necessary to know the exact size of the blockade (as long as it can be distinguished from noise). Thus this method answers the simpler question “Is an AA present or not?” than “Which AA is it?”. This is a binary measurement that converts AA detection from a high precision analog process to a low precision digital one. (In the other 19 channels there is no blockade because there is no freed AA.) See the Supplement to [34] for diagrams showing example flow traces. By successively cleaving the terminal residue of a peptide and using this method in conjunction with the slowdown method introduced here peptide sequencing can be converted into a robust digital measurement process. (Compare this with the analog measurements that lie at the core of existing as well as recently reported methods of protein sequencing; see [34] for the related references.)
6. In some cases the pH values required for achieving slowdown are on the high side (> 10), such high values are known to impact molecular structure. The pH values used for nucleotides and the oligonucleotides is around the physiological value of 7, so the problem of structural change does not apply. With single AAs, a shape change, if it occurs at all, may not be a significant obstacle to identification, in light of Item 5 above. With short peptides (∼8-10 residues) if the change is in the form of stretching, it may actually be beneficial. Thus with the slowdown procedure given here the resulting low bandwidth measurements of translocating short peptides may yield useful sequence information in the form of blockade volumes for them. For example in [35] it is shown that the volumes of short subsequences (as represented by the corresponding blockade current levels) can be used to identify almost all of the proteins in the yeast proteome.
7. The attraction of solution counterions to the analyte may reduce the charge carried by the latter, and hence its mobility, by up to 25% [1].
8. An analyte has to enter the pore within a reasonable amount of time. In [36] a nanopipette-based solution is given; the same effect can be approximated by a tapered *cis* chamber (Figure S-2, Supplement). In the present work hydraulic pressure [37] has been used in all the simulations; see Supplement. However there is a tradeoff: hydraulic pressure also prevents the flow of ions from *trans* to *cis*, thereby effectively halving the pore current.
9. For a brief overview of recent work on the role played by EO in nanopore sensing models see Section S-6 in the Supplement. While the effect of EO in nanopore modeling cannot be discounted entirely, the opposing effect it has on analyte translocation control based on electrophoresis can be nullified in different ways. Thus:
  a. When proteins are sheathed in SDS (sodium dodecyl sulfate) translocation is determined solely by the electrophoretic force, EO is virtually absent [29];
  b. A lipid coating applied to the pore wall can neutralize surface charges on the wall [31], drastically reducing the surface charge density of the pore wall and EO flow in the pore to near zero. This can be used regardless of the analyte so it applies equally to free monomers (nucleotides, amino acids) and oligomers (oligonucleotides and oligopeptides).

## Supporting information

Supplementary information file includes the following material: Properties of analytes (nucleotides, amino acids, oligonucleotides, oligopeptides), including calculation of mobilities and diffusion coefficients; Table of mobility data for all 20 amino acids; Formulation and analysis of Fokker-Planck model; Simulation details; Notes on electroosmotic flow; Additional references.

