## Supplementary File for "Slowing down an analyte in a nanopore"

G. Sampath

---

S-1 Physical and chemical properties of nucleotides, amino acids, oligonucleotides, and oligopeptides
S-2 Schematic of e-cell and tandem pore in three segments
S-3 Fokker-Planck model of translocation of an analyte through a nanopore
S-4 Simulation of e-cell
S-5 Reduction in pore current due to bi-level voltage profiles
S-6 Notes on electroosmotic flow and its role in nanopore models of sensing
Additional References

There is some overlap with the main text to improve readability. Reference numbers 1 through 37 refer to the main text; additional references in this Supplement start at 38 and end at 45.

---

### S-1 Physical and chemical properties of nucleotides, amino acids, oligonucleotides, and oligopeptides

The diffusion constant of a molecule D and its electrophoretic mobility in an electrical field μ are given by the following equations:

$$D = k_B T_R / 6\pi\eta R_{hyd} \qquad \mu = C_{mult} q / 6\pi\eta R_{hyd} \qquad \text{(S-1 a,b)}$$

with $k_B$ = Boltzmann constant ($1.3806 \times 10^{-23}$ J/K), $T_R$ = room temperature (298° K), η = solvent viscosity (0.001 Pa.s for water), $R_{hyd}$ = hydrodynamic radius of the analyte molecule (Å), q = electron charge ($-1.619 \times 10^{-19}$ coulomb), and $C_{mult}$ = charge multiplier for the analyte. With nucleotides $C_{mult}$ is usually taken to be -1.0; with AAs, it depends on the solution pH and is calculated with the Henderson-Hasselbalch equation [30].

*Nucleotides*

The diffusion coefficient and mobility values of the four nucleotides in DNA are given in Table S-1. They are computed from Equations S-1a and b. Hydrodynamic radius values are calculated from an ellipsoid of length 7 Å (= length of stretched mononucleotide, same for all 4 types) with cross-section area calculated from the volume in column 2. Volumes are from [38].

**Table S-1** Nucleotide properties

| Nucleotide | Volume ($10^{-30}$ m$^3$) | Hydrodynamic radius ($10^{-10}$ m) | Diffusion coefficient ($10^{-10}$ m$^2$/s) | Mobility ($10^{-8}$ m$^2$/Vs) |
|---|---|---|---|---|
| dAMP | 349.0 | 4.8790 | 4.4734 | 1.7419 |
| dTMP | 339.0 | 4.8086 | 4.5389 | 1.7674 |
| dCMP | 324.0 | 4.7010 | 4.6428 | 1.8078 |
| dGMP | 359.0 | 4.9484 | 4.4107 | 1.7175 |

*Oligonucleotides*

The diffusion coefficient and mobility of oligonucleotides can be obtained from the individual hydrodynamic radii. The latter are calculated approximately by assuming these analytes to be rigid rods of beads, where a bead corresponds to a nucleotide. The calculations are summarized below.

1) For a rigid rod of length L and radius R, if the motion is limited to the axis of the rod, with no lateral motion, the hydrodynamic radius is given by

$$R_{H.rod} = (LR^2/4)^{1/3} \qquad \text{(S-2)}$$

2) The length $L_{on}$ of an oligonucleotide with k nucleotides $N_1\ N_2\ ...\ N_k$ is approximated as

$$L_{on} = \Sigma_{i = 1, \ldots, K} \; 2R_{H.Ni} \tag{S-3}$$

where $R_{H.Ni}$ = hydrodynamic radius of $N_i$.
3) The radius $R_{on.rod}$ of the rigid rod representing the oligonucleotide is taken to be the average over the hydrodynamic radii of all the nucleotides in the oligo:

$$R_{on.rod} = (1/k)\, \Sigma_{i = 1, \ldots, k}\, R_{H.Ni} \tag{S-4}$$

The calculated values of the hydrodynamic radii and the corresponding diffusion coefficient (using Equation S-1) of the two sample oligonucleotides in Table 1 in the main text are given below in Table S-2.

**Table S-2** Hydrodynamic radius of two sample oligonucleotides. All lengths are in Å (= $10^{-10}$ m).

| **Oligonucl** | **Length** | **Avg radius (Å)** | **hydrad (Å)** | **Diffcoeff ($10^{-10}$ m²/s)** | **Mobility ($10^{-8}$ m²/Vs)** |
|---|---|---|---|---|---|
| AAAAAAAAAA | 9.76 | 4.88 | 8.34 | 2.62 | 10.19 |
| CCAGTTTATA | 9.64 | 4.82 | 8.24 | 2.65 | 10.31 |

*Amino acids*
Write the set of amino acids as

$$\mathbf{AA} = \{A,C,D,E,F,G,H,I,K,L,M,N,P,Q,R,S,T,V,W,Y\} \tag{S-5}$$

The hydrodynamic radius $R_{aa}$ and diffusion coefficient $D_{aa}$ values of the 20 amino acids are given in Table S-3. $R_{aa}$ values are from [39], $D_{aa}$ values are calculated from Equation S-1 a.

**Table S-3** Hydrodynamic radius and diffusion coefficient of the 20 proteinogenic amino acids. $R_{H.aa}$ values are × $10^{-10}$ m, $D_{aa}$ values are × $10^{-10}$ m²/s

| | **A** | **C** | **D** | **E** | **F** | **G** | **H** | **I** | **K** | **L** | **M** | **N** | **P** | **Q** | **R** | **S** | **T** | **V** | **W** | **Y** |
|---|---|---|---|---|---|---|---|---|---|---|---|---|---|---|---|---|---|---|---|---|
| $R_{H.aa}$ | 2.66 | 2.86 | 3.02 | 3.14 | 3.35 | 2.32 | 3.49 | 3.24 | 3.69 | 3.39 | 3.08 | 2.98 | 2.68 | 3.23 | 3.6 | 2.76 | 3.04 | 3.32 | 3.5 | 3.57 |
| $D_{aa}$ | 8.21 | 7.63 | 7.23 | 6.95 | 6.52 | 9.41 | 6.25 | 6.74 | 5.91 | 6.44 | 7.09 | 7.32 | 8.14 | 6.76 | 6.06 | 7.91 | 7.18 | 6.57 | 6.24 | 6.11 |

Following [40] a free AA can be informally written as COOH – Residue – $NH_3$. The charges on the COOH and $NH_3$ components can be calculated from the $pK_1$ and $pK_2$ values for the AA at the respective end; all 20 AAs have $pK_1$ and $pK_2$ values. The residue component has a charge only if the residue has a charged side chain; the corresponding charge can be calculated from the AA's $pK_3$ value and is positive (negative) for positively (negatively) charged side chains (respectively {K, R, H} and {C, D, E, Y}). Table S-4 lists the $pK_1$, $pK_2$, and applicable $pK_3$ values for the 20 AAs, the values are from [30].

**Table S-4** Table of pK values (blank entry = side chain of corresponding AA does not carry a charge)

| **AA** | **A** | **C** | **D** | **E** | **F** | **G** | **H** | **I** | **K** | **L** | **M** | **N** | **P** | **Q** | **R** | **S** | **T** | **V** | **W** | **Y** |
|---|---|---|---|---|---|---|---|---|---|---|---|---|---|---|---|---|---|---|---|---|
| $pK_1$ (--COOH) | 2.34 | 1.96 | 1.88 | 2.19 | 1.83 | 2.34 | 1.82 | 2.36 | 2.18 | 2.36 | 2.28 | 2.02 | 1.99 | 2.17 | 2.17 | 2.21 | 2.11 | 2.32 | 2.38 | 2.2 |
| $pK_2$ (--$NH_3$) | 9.69 | 10.28 | 9.6 | 9.67 | 9.13 | 9.6 | 9.17 | 9.68 | 8.95 | 9.6 | 9.21 | 8.8 | 10.96 | 9.13 | 9.04 | 9.15 | 9.62 | 9.62 | 9.39 | 9.11 |
| $pK_3$ | | 8.18 | 3.65 | 4.25 | | | 6 | | 10.53 | | | | | | 12.48 | | | | | 10.07 |

Thus one can write

$$C_{mult}(AA) = C_{mult\text{-}COOH}(AA) + C_{mult\text{-}NH3}(AA)\,; \qquad AA \in \{A, F, G, H, I, L, M, N, P, Q, S, T, V, W\} \tag{S-6a}$$

$$= C_{mult\text{-}COOH}(AA) + C_{mult\text{-}NH3}(AA) + C_{mult\text{-}res}\,; \qquad AA \in \{C, D, E, H, K, R, Y\} \tag{S-6b}$$

Let the solution pH be p. At this pH, the three $C_{mult}$s are given by

$$C_{mult\text{-}COOH}(AA) = -10^{p} / (10^{p} + 10^{pk1}) \qquad AA \in \{A,C,D,E,F,G,H,I,K,L,M,N,P,Q,R,S,T,V,W,Y\} \tag{S-7a}$$

$$C_{mult\text{-}NH3}(AA) = 10^{pK2} / (10^{pK2} + 10^{p}) \qquad AA \in \{A,C,D,E,F,G,H,I,K,L,M,N,P,Q,R,S,T,V,W,Y\} \tag{S-7b}$$

$C_{mult—res}(AA) = 10^{pK3} / (10^{p} + 10^{pK3})$ AA ∈ {H,K,R} (positively charged AAs) (S-7c)

$= -10^{p} / ( 10^{p} + 10^{pK3})$ AA ∈ {C,D,E,Y} (negatively charged AAs) (S-7d)

The mobility values of the 20 amino acids are then calculated from Equations S-1 b.

**Table S-5** Mobility table for the 20 proteinogenic amino acids. All mobility values are × $10^{-8}$ $m^2/Vs$.

| AA<br>pH | A | C | D | E | F | G | H | I | K | L | M | N | P | Q | R | S | T | V | W | Y |
|---|---|---|---|---|---|---|---|---|---|---|---|---|---|---|---|---|---|---|---|---|
| 0 | 3.180 | 2.939 | 2.777 | 2.689 | 2.500 | 3.647 | 4.834 | 2.612 | 4.591 | 2.496 | 2.745 | 2.825 | 3.139 | 2.614 | 4.706 | 3.060 | 2.774 | 2.548 | 2.418 | 2.366 |
| 1 | 3.055 | 2.678 | 2.480 | 2.541 | 2.210 | 3.503 | 4.550 | 2.513 | 4.464 | 2.402 | 2.622 | 2.603 | 2.877 | 2.465 | 4.572 | 2.900 | 2.594 | 2.443 | 2.331 | 2.239 |
| 2 | 2.193 | 1.417 | 1.152 | 1.630 | 1.023 | 2.514 | 3.404 | 1.826 | 3.690 | 1.745 | 1.810 | 1.459 | 1.567 | 1.570 | 3.769 | 1.905 | 1.574 | 1.731 | 1.714 | 1.460 |
| 3 | 0.574 | 0.248 | -0.316 | 0.219 | 0.161 | 0.658 | 2.584 | 0.489 | 2.606 | 0.467 | 0.442 | 0.270 | 0.282 | 0.339 | 2.665 | 0.430 | 0.319 | 0.442 | 0.470 | 0.326 |
| 4 | 0.068 | 0.027 | -1.924 | -0.933 | 0.017 | 0.078 | 2.427 | 0.059 | 2.338 | 0.056 | 0.052 | 0.030 | 0.031 | 0.038 | 2.395 | 0.049 | 0.036 | 0.052 | 0.057 | 0.037 |
| 5 | 0.007 | 0.001 | -2.692 | -2.294 | 0.002 | 0.008 | 2.215 | 0.006 | 2.306 | 0.006 | 0.005 | 0.003 | 0.003 | 0.004 | 2.364 | 0.005 | 0.004 | 0.005 | 0.006 | 0.004 |
| 6 | 0.000 | -0.019 | -2.802 | -2.659 | -0.002 | 0.000 | 1.216 | 0.000 | 2.301 | 0.000 | -0.001 | -0.004 | 0.000 | -0.002 | 2.359 | -0.002 | 0.000 | 0.000 | 0.000 | -0.002 |
| 7 | -0.006 | -0.186 | -2.820 | -2.708 | -0.019 | -0.009 | 0.205 | -0.005 | 2.277 | -0.006 | -0.017 | -0.045 | 0.000 | -0.019 | 2.339 | -0.022 | -0.007 | -0.006 | -0.010 | -0.021 |
| 8 | -0.064 | -1.198 | -2.883 | -2.763 | -0.175 | -0.090 | -0.130 | -0.054 | 2.064 | -0.061 | -0.160 | -0.390 | -0.004 | -0.182 | 2.163 | -0.204 | -0.066 | -0.060 | -0.095 | -0.195 |
| 9 | -0.542 | -2.729 | -3.379 | -3.183 | -1.080 | -0.735 | -0.980 | -0.453 | 1.019 | -0.503 | -1.052 | -1.749 | -0.034 | -1.120 | 1.234 | -1.276 | -0.541 | -0.495 | -0.703 | -1.257 |
| 10 | -2.145 | -3.950 | -4.827 | -4.551 | -2.235 | -2.620 | -2.121 | -1.774 | -0.336 | -1.793 | -2.374 | -2.683 | -0.313 | -2.318 | 0.226 | -2.698 | -1.973 | -1.807 | -1.950 | -3.299 |
| 11 | -3.046 | -5.463 | -5.521 | -5.292 | -2.503 | -3.523 | -2.400 | -2.503 | -1.700 | -2.411 | -2.715 | -2.834 | -1.659 | -2.596 | -0.050 | -3.036 | -2.684 | -2.457 | -2.370 | -4.514 |
| 12 | -3.179 | -5.887 | -5.617 | -5.401 | -2.534 | -3.649 | -2.432 | -2.611 | -2.226 | -2.497 | -2.755 | -2.850 | -2.906 | -2.628 | -0.585 | -3.075 | -2.784 | -2.549 | -2.422 | -4.735 |
| 13 | -3.193 | -5.937 | -5.627 | -5.412 | -2.537 | -3.662 | -2.435 | -2.622 | -2.295 | -2.506 | -2.759 | -2.852 | -3.143 | -2.631 | -1.813 | -3.079 | -2.794 | -2.559 | -2.428 | -4.758 |
| 14 | -3.195 | -5.943 | -5.628 | -5.413 | -2.537 | -3.663 | -2.435 | -2.623 | -2.302 | -2.507 | -2.759 | -2.852 | -3.168 | -2.631 | -2.292 | -3.079 | -2.796 | -2.560 | -2.428 | -4.761 |

*Peptides*

Let peptide P have k (> 1) residues in the sequence $A_0$ $A_1$ ... $A_{k-2}$ $A_{k-1}$ (written from C-terminal to N-terminal). As with single free AAs the oligopeptide of P can be written informally as

$$\mathbf{P} = COOH – A_0 - A_1 - ...- A_{k-2} - A_{k-1} – NH_3 \quad \text{(S-8)}$$

with - between residues representing the peptide bond

For a given pH the charge multiplier $C_{mult-pep}$ for **P** is

$$C_{mult-pep} = C_{mult-COOH} (A_0) + C_{mult-res} (A_0) + ... + C_{mult-res} (A_{k-1}) + C_{mult-NH3} (A_{k-1}). \quad \text{(S-9)}$$

The values of $C_{mult-pep}$ and the corresponding mobilities for the five example peptides used in this study are given in Table S-6 for different pH values. As noted in [40], this assumes that there are no interactions between residues, which would be true of fully stretched peptides.

As an example, the $C_{mult}$ value for the analyte IVHSMSWALP at a given pH value is

$$C_{mult-IVHSMSWAL} = C_{mult-COOH} (I) + C_{mult-res} (V) + C_{mult-res} (H) + ... + C_{mult-res} (L) + C_{mult-res} (P) + C_{mult-NH3} (P)$$
$$= C_{mult-COOH} (I) + 0 + C_{mult-res} (H) + 0 + ... + 0 + C_{mult-NH3} (P) \quad \text{(S-9a)}$$

As with oligonucleotides, oligopeptides are considered to be rigid rods of beads, where a bead corresponds to an amino acid (internal residue). The hydrodynamic radius of the oligopeptide modeled as a rod of length $L = L_{op}$ and R = radius $R_{op}$ is given by Equation S-2.

The length $L_{op}$ of an oligopeptide with k amino acids (residues) $A_1$ $A_2$ ...$A_k$ is approximated as

$$L_{op} = \Sigma_{i = 1, ..., k} 2R_{H.Ai} - (k-1) \delta v \quad \text{(S-10)}$$

where δv is the van der Waals volume of water (= 0.0186 $nm^3$) [41].

Once again the radius $R_{op}$ of the rigid rod representing the oligopeptide is taken to be the average over the hydrodynamic radii of all the amino acids in the oligo:

$$R_{op} = (1/k) \Sigma_{i = 1 , ..., k} R_{H.Ai} \quad \text{(S-11)}$$

The calculated values of $R_{H.op}$, the hydrodynamic radius of the oligopeptide, for the five sample oligopeptides in Table 1 in the main text and the corresponding diffusion coefficients (based on Equation S-1) are given below in Table S-6.

**Table S-6** Diffusion coefficient and mobility of five sample oligopeptides

| Oligopeptide | Length (Å) | Avg radius (Å) | pH | Hydrodynamic radius $R_{H.op}$ (Å) | Diffusion coefficient ($10^{-10}$ m$^2$/s) | $C_{mult}$ | Mobility ($10^{-8}$ m$^2$/Vs) |
|---|---|---|---|---|---|---|---|
| RRRRRRRRRR | 72.00 | 3.60 | 7 | 6.15 | 3.55 | 8.00 | 11.04 |
| HWVEDVDLTP | 63.84 | 3.19 | 7 | 5.46 | 4.00 | -3.00 | -4.67 |
| IVHSMSWALP | 61.76 | 3.09 | 7 | 5.28 | 4.13 | 0.09 | 0.14 |
| IVFSMSWALP | 61.48 | 3.07 | 7 | 5.26 | 4.15 | 0.00 | 0.00 |
| IVFSMSWALP | 61.48 | 3.07 | 10 | 5.26 | 4.15 | -0.72 | -1.16 |
| RRRRRIVFSM SWALP | 97.48 | 3.25 | 7 | 6.36 | 3.43 | 3.99 | 5.33 |

### S-2 Schematic of e-cell and tandem pore in three segments

Figure S-1 shows a basic e-cell with a single nanopore, as well as an extended e-cell with a tandem pore in 3 segments corresponding to the positive voltage segment, the conducting layer for use as an electrical terminal, and the negative voltage segment. This figure is a repeat of Figure 1 in the main text.

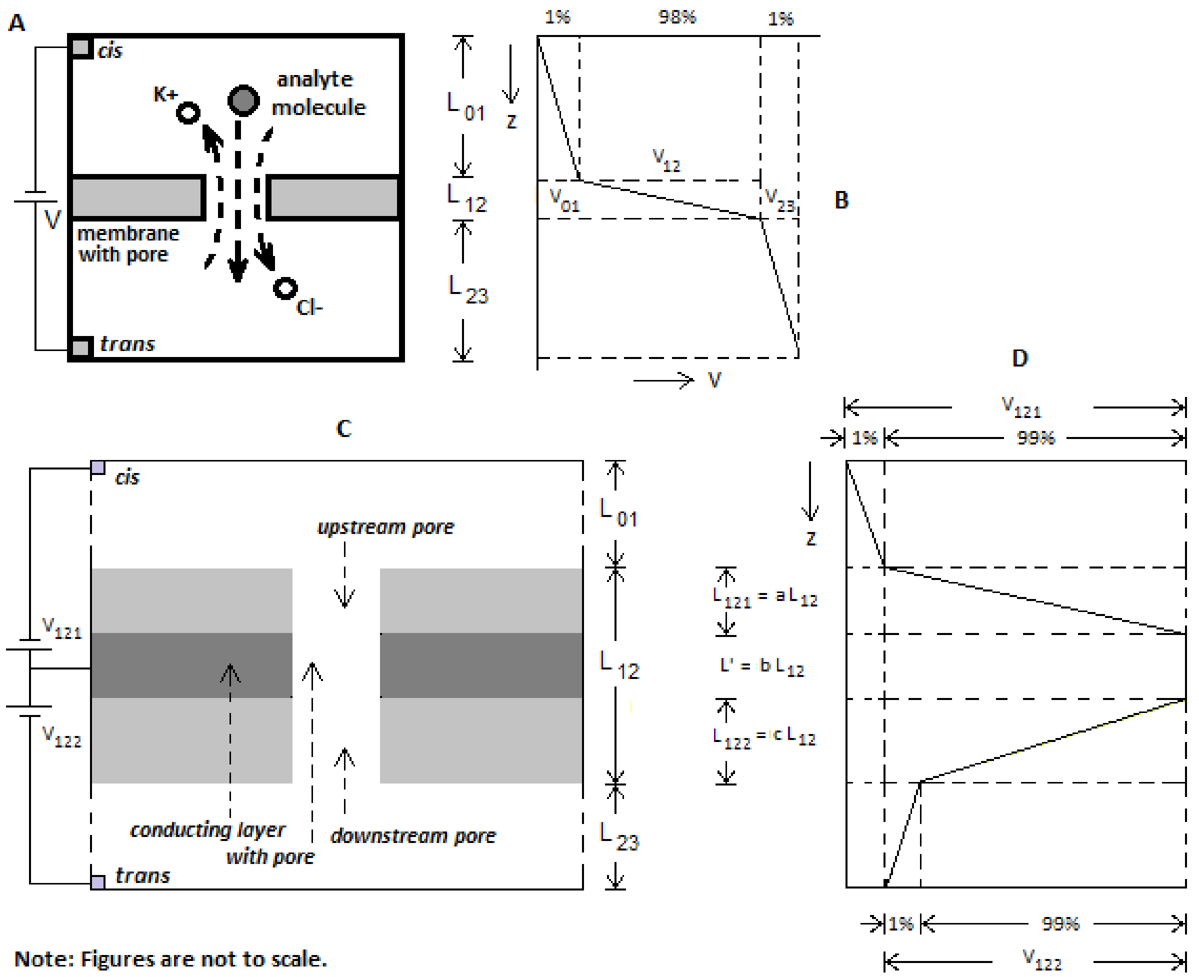


**Figure S-1**. Schematic of electrolytic cell (e-cell); membrane containing nanopore separates *cis* and *trans* chambers containing salt (KCl) solution. (a) Ionic current flow in e-cell is due to $K^+$ and $Cl^-$ ions; analyte translocating through pore causes reduction of open pore current; (b) 98% of voltage V applied between *cis* and *trans* drops across pore; (c) Two pores in tandem with different voltages across each with intervening conducting membrane layer; (d) Positive voltage $V_{121}$ across upstream pore of length $L_{121} = aL_{12}$, with 99% dropping across pore, the other 1% across *cis*; conducting membrane layer at constant potential over thickness $bL_{12}$; negative voltage $V_{122}$ across downstream pore of length $L_{122} = cL_{12}$, with 99%

dropping across pore, the other 1% across *trans*. In simulations $L_{12}$ = 11 nm, a = c = 5/11, b = 1/11. Voltages $V_{121}$ and $V_{122}$ are varied to give different profiles so that analyte can be trapped for long durations; their values depend on analyte and pH of solution in e-cell, see Figure 2 in main text.

**S-3 Fokker-Planck model of translocation of an analyte through a nanopore**
The following is an abbreviated treatment of the model, the extended analysis can be found in the Supplement to [13].

The probability function for the analyte trajectory passing through the pore is G(z,t), which is defined by:

$$\partial G/\partial t + v_z\, \partial G/\partial z = D\, \partial^2 G/\partial z^2, \qquad z \in [0, L=L_{12}] \qquad \text{(S-12)}$$

where $v_z$ is the drift velocity given by

$$v_z = \mu V_{12}/L \qquad \text{(S-13)}$$

The following initial and boundary value conditions apply:

1) The particle is released at z = 0 and time t = 0:

$$G(0,t=0) = \delta(z) \qquad \text{(S-14 a)}$$

2) The particle is captured at z = L:

$$G(L,t) = 0 \qquad \text{(S-14 b)}$$

3) The particle is reflected at z = 0:

$$D\, \partial G(z,t)/\partial z\, |_{z=0} = v_z\, G(z,t) \qquad \text{(S-14 c)}$$

Following the analysis in the Supplement to [13], the statistics of translocation of an analyte through the nanopore are as follows:
1) The mean E(T) is

$$E(T) = (L^2/D\alpha)[1 - (1/\alpha)\,(1 - \exp(-\alpha))] \qquad \text{(S-15)}$$

2) The variance $\sigma^2(T)$ is

$$\sigma^2(T) = (L^2/D\alpha^2)^2\, (2\alpha + 4\alpha \exp(-\alpha) - 5 + 4 \exp(-\alpha) + \exp(-2\alpha)) \qquad \text{(S-16)}$$

where σ is the standard deviation.
3) With pure diffusion in the absence of an electrophoretic field the drift velocity $v_z = 0$; the mean and variance are given by

$$E_0(T) = L^2/2D \qquad \text{(S-17)}$$
$$\sigma_0^2(T) = (1/6)\,(L^4/D^2) \qquad \text{(S-18)}$$

**S-4 Simulation of e-cell**
The structure of the basic e-cell used in the simulation is shown in Figure S-2. The taper in the *cis* chamber aids translocation of an analyte released at the top of the chamber to the pore entrance. Additionally hydraulic pressure can be used to ensure delivery of an analyte from the top of the *cis* chamber to the pore entrance; ΔP is in the range 1 to 2.5 atm.

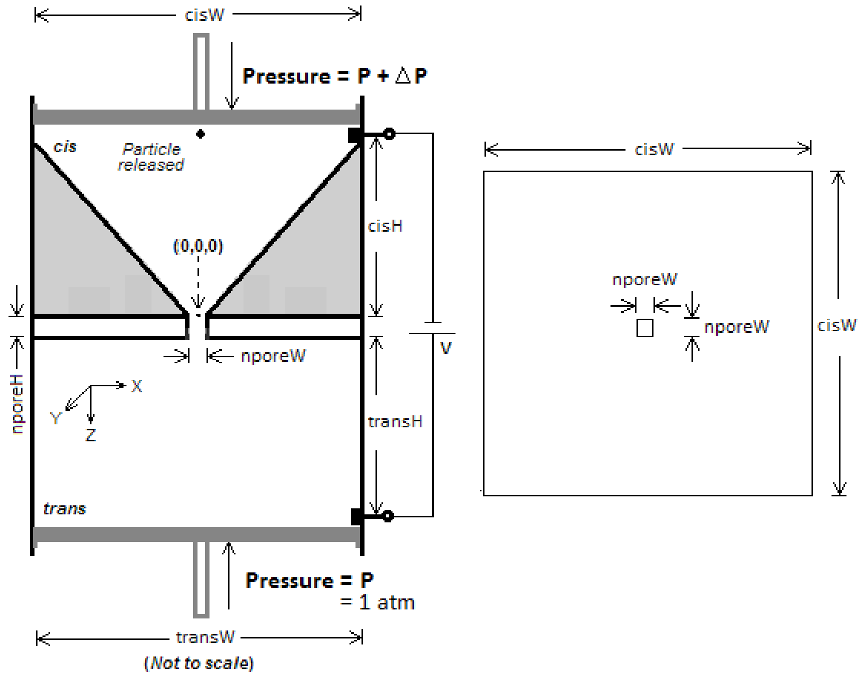


**Fig. S-2** Geometry (symmetric in X and Y) of basic e-cell. Structure shown in plan on the right. This geometry is carried over to the e-cell in Figure S-1 (same as Figure 1 in the main text)

*Simplifying assumptions for the simulation*

1) A free nucleotide or amino acid is a dimensionless particle that is reflected off the walls of *cis*, nanopore, and *trans*. Here 'nanopore' is the three-level structure in Figure S-1.

2) The hydraulic pressure field superimposed on diffusion by the piston is assumed to result in Poiseuille flow in all three channels: *cis*, pore, and *trans*. The hydrodynamic velocity $v_{hyd}$ of a particle under Poiseuille flow is given by

$$v_{hyd} = (P_{atm} + \Delta P)\, R^2_{channel} / 8\, \eta\, H_{channel} \qquad \text{(S-19)}$$

where $P_{atm}$ = atmospheric pressure, $\Delta P$ = applied pressure on piston in *cis*, $R_{channel}$ is the radius of the channel, $\eta$ is the viscosity of the solution, and $H_{channel}$ is the height of the channel.

3) The particle moves in the pore under the influence of electrophoresis and diffusion (and hydraulic pressure, whose effect inside the narrow pore is minor). Electroosmotic flow is excluded.

3) The displacements due to hydraulic pressure and electrophoresis are always in the z direction.

*Calculations*

1) The electric field across a pore of interest is $E = V/L$, where V is the applied voltage across the pore and L is the thickness (height) of the pore. The electrophoretic drift velocity is $v_z = \mu E$ where $\mu$ is the mobility and $E = V_{12}/L$ is the electric field.

2) The displacement of a particle inside a pore is the vector sum of the displacements due to diffusion, electrophoresis, and hydraulic pressure. Diffusive movement occurs in a random direction in 3-d space and is simulated as a displacement during a small time interval $\Delta t$ (set to the simulation time step of 1 picosecond = $10^{-12}$ s). The diffusion displacement size is $\sqrt{(6D\Delta t)}$, where D is the diffusion constant of the analyte. A random direction is generated as a unit vector on the surface of a unit sphere. This vector is then multiplied by the diffusion step size and the translation due to drift in the direction of the electric field and translation due to hydraulic pressure are added to it. The resulting position is accepted if it is inside *cis*, pore, or *trans*, and rejected otherwise.

3) The simulation is stopped when the total number of steps reaches $2 \times 10^9$ or when the particle crosses into *trans*.

*Constants used*

1) Atmospheric pressure $P_{atm}$ = 106803 Pa (= pressure unit Pascal)

2) Viscosity of solution 0.001 Pa.sec

*Variables recorded*

1) The *dwell time* of a particle inside the nanopore is the time spent by the particle inside the pore from the time of entry from *cis* to the time of regression into *cis* (or no regression occurs because the maximum number of steps in the simulation has been reached). The *maximum dwell time* is the maximum such time over the full simulation or until the particle has exited to *trans*.
2) The *translocation time* of a particle through a nanopore is the time between the time of entry from *cis* into the pore to the time of exit into *trans* without ever once regressing into *cis*.
3) Simulation *execution time* is the clock time from start of simulation to end of simulation. (This time is recorded but not reported in the results.)

*Simulation program*
Simulation was done with a customized program in C.

**S-5 Reduction in pore current due to bi-level voltage profiles**
The downside to the slowdown method given above is a decrease in the pore ionic current $I_{pore}$. $I_{pore}$ has four components: the currents due to K+ and Cl- ions moving forward from *cis* to *trans* (c→t) and the corresponding reverse currents from *trans* to *cis* (t→c). One can write

$$|I_{pore}| = |I_{Cl-,c\to t} - I_{Cl-,t\to c}| + |I_{K+,t\to c} - I_{K+,c\to t}| \qquad \text{(S-20)}$$

**Table S-7** Increase in translocation times of Cl- ions (*cis → trans*) and K+ (*trans → cis*) with bi-level voltage profile. Profile has the form (positive voltage segment, voltage over conductor layer = 0, negative voltage segment); see Figure S-1. Data for ions flowing in reverse are not shown.

| Voltage profile (V, V, V) | E(T) with voltage profile ($10^{-8}$ s) | | Normal pore voltage (V, V, V) | E(T) with normal pore voltage ($10^{-8}$ s) | |
|---|---|---|---|---|---|
| | Cl- | K+ | | Cl- | K+ |
| (0.5,0,-0.12) | 5.83 | 5.98 | 0.15 | 0.55 | 0.57 |
| (0.5,0,-0.15) | 12.4 | 12.6 | 0.25 | 0.85 | 0.88 |
| (0.5,0,-0.1) | 3.69 | 3.79 | 0.35 | 0.41 | 0.420 |

The decrease can be estimated from the corresponding translocation/dwell times of ions in the pore. Table S-7 shows the increased translocation times due to $Cl^-$ ions flowing from *cis* to *trans*, which results in a net decrease in the current. As the normal $K^+$ ion flow from *cis* to *trans* is small the effect of the voltage profile on it is much smaller. A decrease similar to that with $Cl^-$ going from *cis* to *trans* occurs in the K+ current from *trans* to *cis*. The smaller effects due to $K_{+,c\to t}$ and $Cl_{-,t\to c}$ are not included in the table. Since the current delivered by an ion from *cis* (*trans*) to *trans* (*cis*) is inversely proportional to the translocation time through the pore that separates the two chambers the result is a steep fall in the current to ~10% of that without the opposing voltage.

**S-6 Notes on electroosmotic flow: its role in nanopore models of sensing and analysis**
The role of electroosmotic flow in nanopores due to surface charged residues on the pore wall has been studied extensively [4-7,42-45]. Counterions attracted to these charged residues reduce the ionic flux through the pore, sometimes enough to reverse the direction of the analyte. Analyte translocation is thus subject to the competing forces of electrophoresis (EP), diffusion, and electroosmotic (EO) flow [7]. The balance is also affected by the electrolyte. Thus with KCl for the electrolyte translocation events are dominated by electrophoresis, whereas with low concentrations of LiCl the dominant force is EO. Intermediate concentrations result in EP and EO balancing out so that diffusion becomes the determinant of analyte translocation through the pore [7]. At a salt concentration of roughly 100 mM translocation goes from EO-dominated to EP-dominated [45].
The influence of EO on nanopore-based analysis and sensing can be considered from different perspectives:
1) As it tends to oppose EO it has the beneficial effect of retarding analyte translocation thus reducing the detection bandwidth required. In the extreme case when EO overwhelms EP analyte detection becomes a problem.
2) With proteins EO has the further effect of uncoiling and stretching a folded protein thus exposing the residues in the primary sequence to inspection and discrimination [43].
3) With biological nanopores that are anchored in a fragile bilipid layer, the high salt concentrations required to minimize EO can rupture the bilipid layer [46]. This may not be a problem with synthetic nanopores.

4) In the present context the effect of EO can be more or less nullified:

- Thus when proteins are sheathed in SDS (sodium dodecyl sulfate) to invest them with a uniform charge along their length translocation is determined by the electrophoretic force, EO is virtually absent [29].
- A lipid coating applied to pore wall can neutralize surface charges on the wall [31], in effect drastically reducing the surface charge density and EO flow to zero.
